# Computational Principles of Auditory Object Formation Are Revealed by Repetition Coherence

**DOI:** 10.64898/2026.01.14.699508

**Authors:** Berfin Bastug, Yue Sun, Federico Adolfi, Erich Schröger, David Poeppel

## Abstract

A foundational task for perceptual systems is to identify objects. In hearing, repetition — for example of a feature critical to object identity — is a powerful cue for object formation. The principles governing how repetition may underpin this process are not well understood, particularly for the noisy, approximate repetition typical of biological environments. To characterize the perceptual organization of acoustic scenes into objects, we adapted the very successful motion-coherence paradigm from vision research, developing a repetition-coherence framework to investigate the evidence accumulation that yields auditory perceptual objects. Participants heard dense tone-cloud sequences in which subsets of tones repeated across cycles. The proportion of repeating tones defined coherence, and the length of the repeating unit was varied. They performed two tasks: repetition detection, which captures the endpoint of object formation, and sensorimotor synchronization, which provides a continuous readout of object formation as it unfolds. The convergence between the two tasks validates sensorimotor synchronization as an online behavioral probe of an otherwise covert process. We discovered that at high coherence both detection and stable tracking were achieved at the same number of cycles, independent of other factors. Yet within this fixed integration regime, longer repeated units were less likely to give rise to a perceptual object, dissociating the likelihood of object emergence from its timing. This scale-independent integration limit suggests that auditory object formation is governed by two interacting principles: extraction of statistical regularities and a fixed integration window. The behavioral signatures parallel known neurophysiological dynamics, providing a helpful link to interpret them.

## Introduction

The ecologically typical acoustic environment contains (often continuous) streams of overlapping sounds that convey information across multiple temporal scales. Individual sounds typically lack clear boundaries, and their components often coincide in time, creating a complex mixture for the auditory system to process. One central challenge is to decompose this input and organize it into meaningful units (auditory objects; e.g., Bizley & Cohen, 2013; Griffiths & Warren, 2004) that can be tracked and interpreted (Bregman, 1994).

One powerful cue for organizing sound is repetition. Repeated structures are common in auditory scenes, from footsteps to rhythmic patterns in speech and music (Keyser, 2025; Margulis, 2013; Poeppel & Assaneo, 2020). Repetition offers a reliable statistical regularity that allows the auditory system to guide perceptual grouping (McDermott et al., 2011; Winkler et al., 2009), enhance predictability (Baldeweg, 2006), and support change detection (Barascud et al., 2016; Chait et al., 2008).

Listeners are highly sensitive to temporal regularities in sound (Agus et al., 2010; Bendixen et al., 2009; Garrido et al., 2013; Luo et al., 2013; Wacongne et al., 2011). This sensitivity has long been investigated, for example, with mismatch negativity (MMN) paradigms, wherein infrequent deviations from a regular tone sequence evoke a rapid neural response (Näätänen et al., 2007; Paavilainen, 2013; Schröger et al., 2014). Besides violations of established regularities, studies have also demonstrated that listeners can detect the emergence of temporal regularities from an initially random sound sequence (Barascud et al., 2016; Heilbron & Chait, 2018; Southwell et al., 2017). For instance, Barascud et al. (2016) presented listeners with sequences of pure tones whose frequencies fluctuated randomly within a specified range, then transitioned into a regular sequence by repeating a short segment of the preceding random sequence. Participants detected these transitions after a single cycle of repeated acoustic material. At the neural level, this perceptual shift elicited a sustained increase in magnetoencephalographic (MEG) responses, in contrast to the transient responses typically observed in MMN paradigms (Näätänen et al., 2007; Paavilainen, 2013). The authors proposed that this sustained response reflects an ongoing process that tracks the repeating structure as it is established.

Barczak et al. (2018) characterized this ongoing process by recording neural activity in the macaque auditory pathway while the animals passively listened to tone sequences like those used by Barascud et al. (2016). They observed that thalamocortical activity gradually aligned with the regular sequence after its onset, with alignment strength increasing over successive repetition cycles before plateauing — a dynamic they interpreted as a candidate mechanism for grouping continuous acoustic input into auditory objects.

Notwithstanding these important advances, critical gaps in our understanding remain. Prior work has typically contrasted fully repeating with fully random sequences (e.g., Barascud et al., 2016; Barczak et al., 2018). But natural auditory regularities are rarely exact and often partial, with relevant patterns embedded within a constantly changing soundscape. Furthermore, previous studies used sequences of perceptually resolvable single tones, which made the formation of an auditory object (i.e., a multi-tone sequence) a concatenation of these events. In contrast, perception of complex auditory signals often requires grouping acoustic features that unfold over time, where individual fragments may not correspond to known auditory events. These limitations leave open a fundamental question: how does the auditory system integrate partial sensory evidence over time to form coherent auditory objects?

To address this question, we adapted an experimental design that has been highly productive in vision: the *motion coherence* paradigm (Britten et al., 1992; Kim & Shadlen, 1999; Shadlen & Kiani, 2013). The paradigm parametrically varies the proportion of coherently moving dots to manipulate signal strength. This graded control reveals the principles by which the brain integrates sensory evidence to judge the global motion of a dot cloud. The same approach, applied to repetition in audition, can expose the principles of sensory evidence integration underlying auditory object formation. We therefore introduce a *repetition-coherence* paradigm. Our stimuli (Figure 1A-C) were tone clouds: clusters of brief tones randomly distributed across a time-frequency grid (Agus & Pressnitzer, 2021). At any moment, ten tones overlap, making individual tones within a time frame difficult to resolve. Thus, detection of a repeating pattern relies on integration across time frames. We manipulated repetition coherence by freezing a proportion of tones across repetition cycles while randomly distributing the remainder. This produced a graded coherence axis from fully random to fully repeating (Figure 1C).

**Figure 1.**
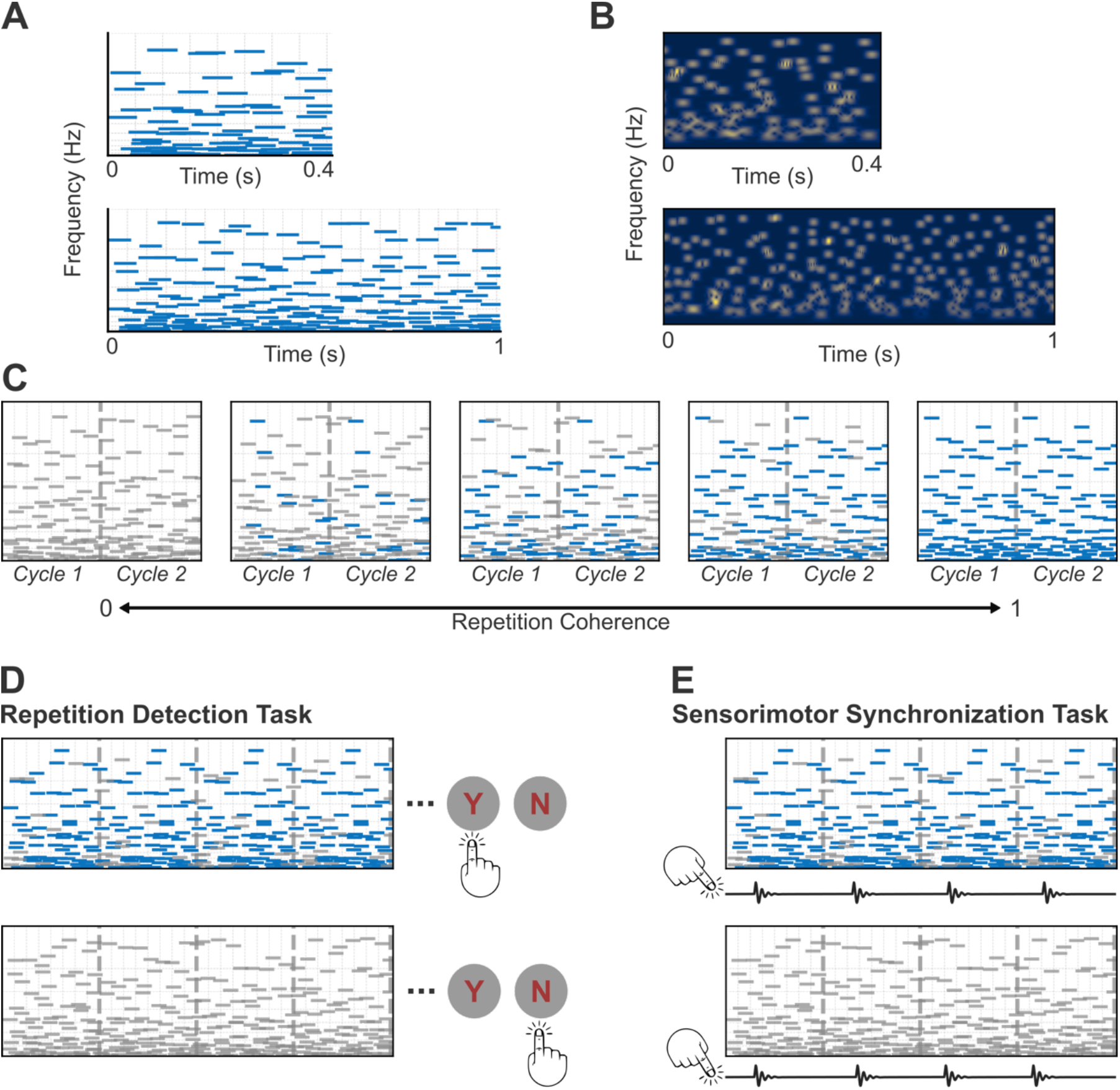
Stimulus construction and task design. Spectrogram (A) and cochleagram (B) representations of tone clouds with the shortest (0.4-s top) and longest (1.0-s bottom) unit durations (cochleagram adapted from Norman-Haignere et al. (2022)). Tone clouds were generated using a time-frequency grid. Frequencies spanned 200–3000 Hz. Each grid column lasted 0.05 s, and cells were populated with brief tones of the same duration. Unit duration determined the number of columns. Three unit durations were used in the experiment (0.4, 0.7, and 1.0 s). (C) Example of 0.4-s tone-cloud units at five coherence levels. Gray lines indicate newly generated tones on each repetition; blue lines indicate frozen tones. From left to right, coherence increases (0, 0.22, 0.44, 0.67, 1), progressing from fully random to fully repeating sequences. (D–E) Experimental tasks. (D) Repetition detection: top, first five repetitions of a 0.78-coherence sequence (“yes” trial); bottom, five repetitions of a 0-coherence sequence (“no” trial). Sequences ended upon participant response. (E) Sensorimotor synchronization: first five of 30 repetitions at coherence levels 0.78 and 0. Participants tapped in synchrony with the perceived repeating pattern.

Participants performed two complementary psychophysical tasks. In a repetition detection task (Figure 1D; cf. Agus et al., 2010; Agus & Pressnitzer, 2021; Guttman & Julesz, 1963), participants pressed a button to indicate whether a repeating pattern was present in the tone-cloud sequences. This task measured the minimum evidence required to perceive a repeating tone pattern as an auditory object and offered a single decision at the endpoint of the integration process. We further asked whether the gradual alignment of neural activity during object formation (Barczak et al., 2018) has a functional role in perception that can be probed behaviorally. To this end, participants also performed a sensorimotor synchronization (SMS) task (Figure 1E; cf. Repp, 2005; Repp & Su, 2013), tapping fingers in synchrony with the perceived repeating pattern. This task tracked how participants’ motor output evolved in response to the repeating pattern throughout the sequence, providing a continuous index of both the initial detection of the pattern and participants’ ability to maintain its representation over time.

Finally, we used an autocorrelation-based observer model as a conceptual test to query repetition-based segmentation, distinguishing aspects of temporal integration that a bottom-up periodicity detector can account for from those requiring additional mechanisms. Altogether, these tools help us characterize the computational principles by which the auditory system reorganizes complex, dense acoustic input into coherent objects.

## Results

### Longer units require more coherence to detect, but not more cycles

Participants listened to long sequences of tone clouds and detected repeating tone patterns at varied repetition-coherence levels: from 0 to 1. Sequences looped continuously until participants pressed a button. This approach measured both the coherence required for detection and the time taken to reach it. We also varied the duration of a tone cloud (i.e., unit duration) to assess whether coherence effects were consistent across unit durations or whether evidence integration depended on the temporal window over which repetition unfolded.

Detection performance increased sigmoidally (Figure 2A), from 15% yes responses at 0 coherence (false alarms) to 96% detection at 1.0 coherence. Examining repetition coherence at different unit durations, we observed distinct psychometric curves (Figure 2B). A two-way repeated-measures ANOVA quantified how coherence and unit duration jointly impact detection accuracy. This analysis showed a significant main effect of coherence (F (2.29, 59.60) = 352.05, p < 0.001, 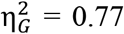) and unit duration (F (1.59, 41.43) = 186.05, p < 0.001, 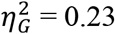), with overall lower accuracy for longer units (Table S1). Importantly, there was a significant interaction between unit duration and coherence (F (6.7, 174.15) = 22.94, p < 0.001, 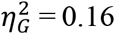), which indicates a scale-dependent influence of coherence on detection performance.

**Figure 2.**
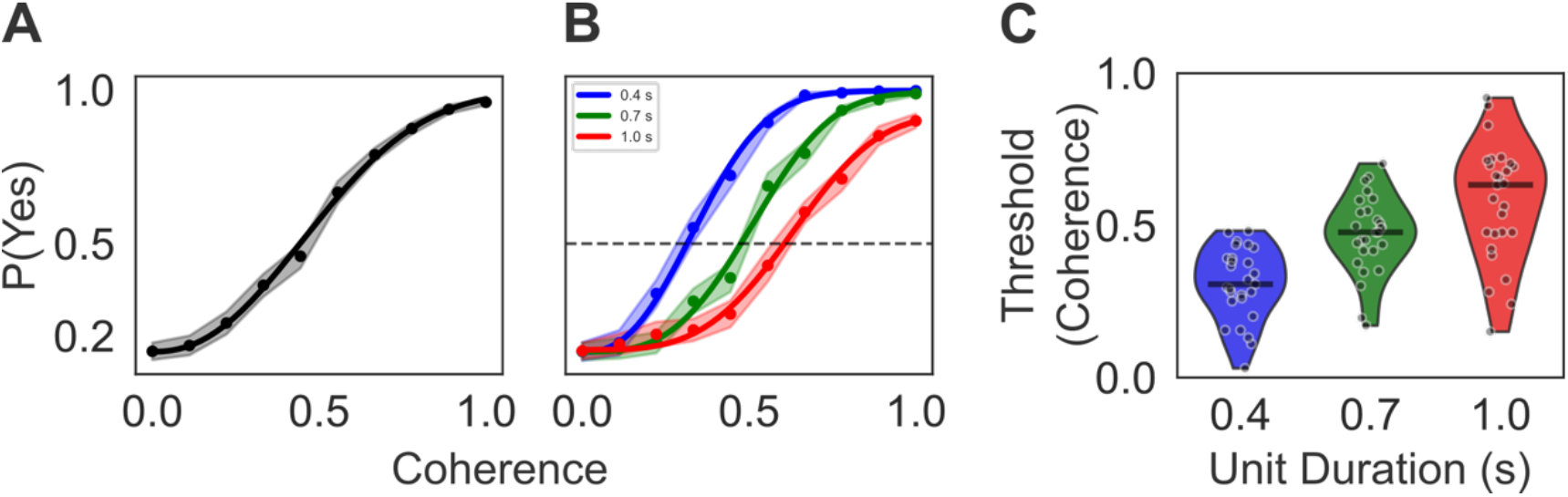
Repetition coherence required for detection. (A-B) Proportion of “yes” responses as a function of repetition coherence. Dots indicate observed data; solid lines show Weibull fits. Shaded areas represent the standard error of the mean across participants. (A) Overall performance averaged across all unit durations. (B) Performance shown separately for each unit duration (0.4 s: blue, 0.7 s: green, 1.0 s: red). The dashed line indicates the 50% ‘yes’ point, taken as the detection threshold. (C) Violin plot of the distribution of threshold parameters (i.e., coherence level at which performance exceeds 50%) from individual Weibull fits for each unit duration.

The performance differences across durations arose from shifts in detection thresholds (defined here as the coherence level required to achieve 50% detection probability). As shown by a repeated-measures ANOVA on Weibull fits, thresholds increased for longer unit duration (Figure 2C). That is, shorter units allowed object detection at significantly lower coherence levels, longer units required stronger coherence to reach comparable performance (Table S2).

Next, we asked how much time participants needed to integrate evidence to detect repetition. We measured reaction times (RTs) as the elapsed time (in seconds) from the onset of the tone-cloud sequence. Detection was fastest for shorter units, then for mid- and then long-duration units (Figure 3A). At low coherence (≤ 0.33), when detection was below chance, RTs were similar across different unit durations. With no robust percept of a repeating pattern, the auditory system treated all durations similarly. The first significant divergence appeared at 0.33 coherence, at which point the 0.4-s units surpassed the detection threshold. At 0.56 coherence, when 0.7-s units reached detection threshold, RTs diverged across all durations (Table S3).

**Figure 3.**
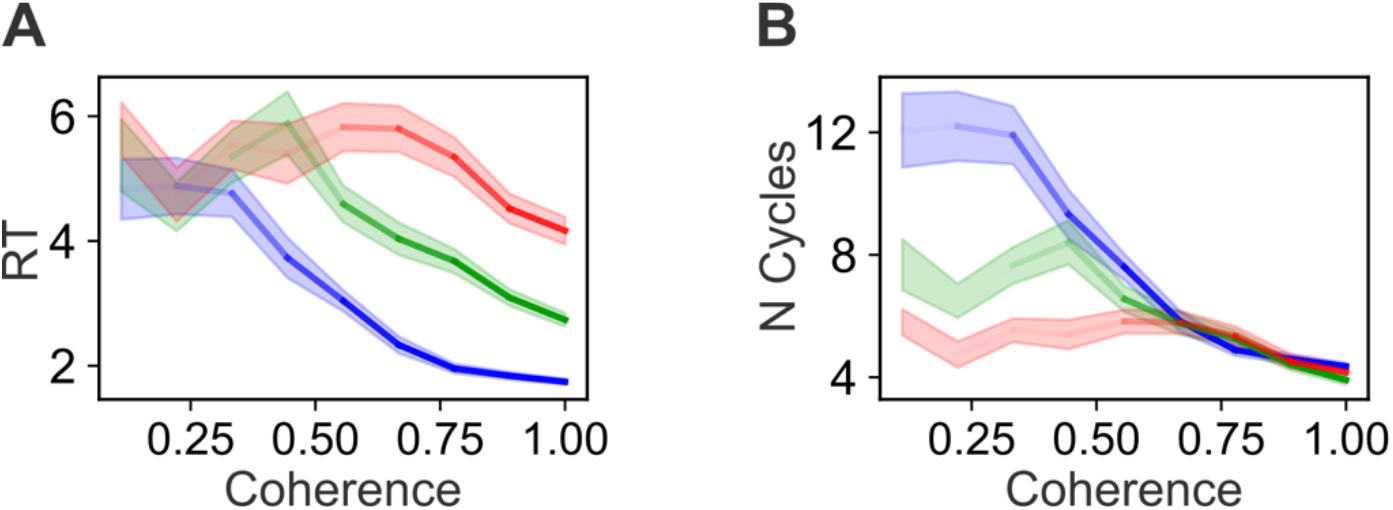
Time needed to integrate sensory evidence for repetition detection. Reaction times and number of cycles required for detection as a function of repetition coherence and unit duration (0.4 s: blue, 0.7 s: green, 1.0 s: red). (A) Group-level mean reaction times (RTs) as a function of repetition coherence and unit duration. Line opacity corresponds to mean accuracy, with darker segments indicating higher proportions of correct responses. Shaded areas indicate ±1 standard error of the mean (SEM). (B) Group-level mean number of cycles required for repetition detection, plotted against repetition coherence for each unit duration (0.4 s: blue, 0.7 s: green, 1.0 s: red). As in panel A, shaded areas show ±1 SEM, and line opacity scales with accuracy.

Recasting RTs from time (seconds) to number of repetition cycles (N cycles) provides a complementary perspective. At low coherence, because RTs were similar across durations, shorter units necessarily sampled more cycles than longer ones within the same elapsed time. As coherence increased, however, N cycles converged across durations (Figure 3B). At 0.67 coherence, cycle counts fully converged. At ceiling coherence, detection was achieved around the fourth cycle after stimulus onset, regardless of unit duration (though a modest difference remained, with 0.7-s units requiring slightly fewer cycles than 0.4-s units; Table S3). This pattern reveals a dissociation within the detection task itself: coherence and unit duration interact to shape detection thresholds, yet once coherence is sufficient for detection, the number of cycles required is effectively scale-independent.

### Sensorimotor synchronization mirrors detection across coherence but not across unit duration

Having established these detection signatures, we examined whether the gradual neural alignment to repeating structure reported by Barczak et al. (2018) is reflected in a corresponding behavioral dynamic, one that can be tracked through sensorimotor synchronization. Participants began tapping from trial onset and continued through 30 cycles, providing a continuous readout that captured both the initial search for the repeating structure and the subsequent tracking of the detected pattern.

First, we assessed how faithfully participants tracked the repeated pattern at different coherence levels. We classified a trial as successfully synchronized when the distribution of tapping phases was more consistent than expected by chance (see Materials and Methods for details). The proportion of successfully synchronized trials increased sigmoidally with coherence, which mirrored the pattern observed in the detection task (Figure 4A). Two fundamentally different readouts, a single perceptual judgement and a continuous motor output, yielded converging psychometric functions.

**Figure 4.**
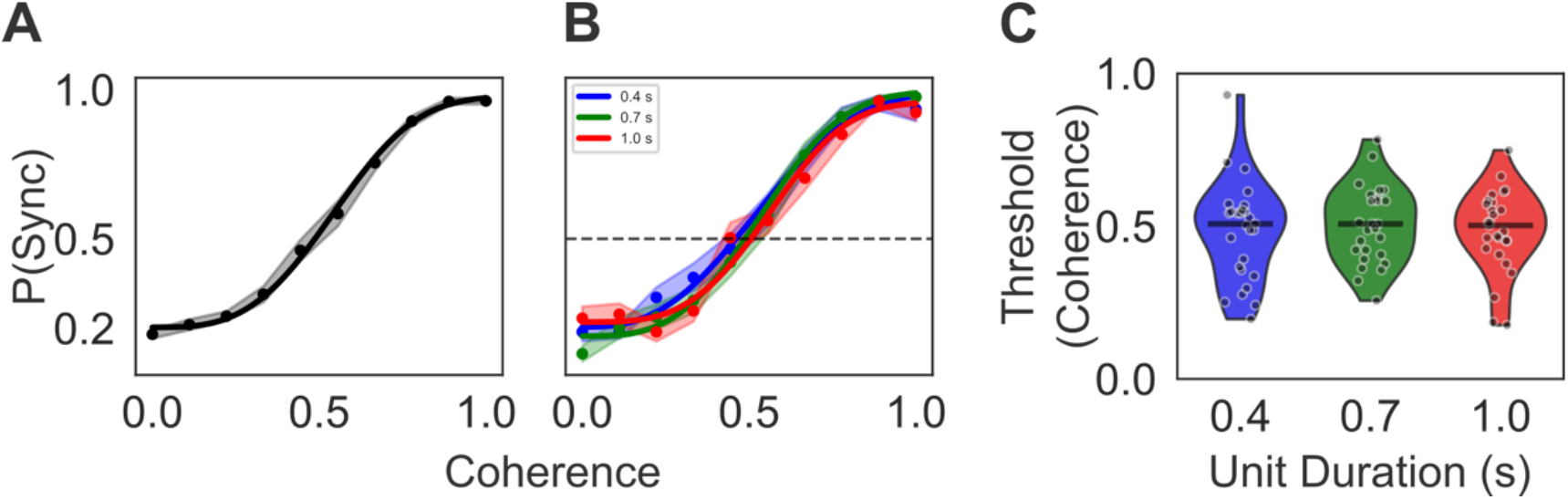
Results of the sensorimotor synchronization (SMS) task. Proportion of trials with successful synchronization is plotted as a function of repetition coherence. (A) Overall performance averaged across unit durations. (B) Performance shown separately for each unit duration (0.4 s: blue, 0.7 s: green, 1.0 s: red). Dots represent observed data; solid lines show Weibull fits, with shaded regions showing the standard error of the mean across participants. The dashed line indicates 50% of trials showing successful synchronization, taken as the threshold for the SMS task. (C) Distribution of threshold parameters from individual Weibull fits for each unit duration (color coding as in B).

We then investigated whether unit duration modulated synchronization performance as it did in detection. Unlike the distinct psychometric curves observed in detection (Figure 2B), synchronization performance remained uniform across durations (Figure 4B), with comparable performance for 0.4-s, 0.7-s, and 1.0-s units (Table S4). Similarly, threshold coherence values estimated from Weibull fits did not differ across durations (Figure 4C; Table S2). Synchronization performance was thus insensitive to the factor that shifted detection thresholds.

Next, to examine the dynamics at a higher resolution, we zoomed in on participants’ within-trial synchronization performance to quantify how it evolves over the course of a trial. To this end, we applied a sliding-window analysis. Each window spanned 10 repetition cycles (e.g., 4 s for a 0.4-s unit) and advanced by one cycle. Within each window, we quantified tapping phase consistency as the z-score of the mean resultant length against a null distribution of random phases (see Materials and Methods). At low coherence (≤ 0.33), tapping phase consistency within each window was statistically indistinguishable from a random phase distribution (ps > 0.05; Figure 5A, values below the dashed baseline), i.e., taps were spread across all possible phases. As coherence increased, tapping phase consistency within each window gradually rose over the course of the trial. Initially dispersed tapping phases progressively converged to a stable position, thereby becoming significantly distinguishable from a random phase distribution. At the highest coherence level, this evolution followed a two-stage trajectory: a rapid build-up, followed by a stabilization during which performance plateaued (Figure 5A). This buildup-and-plateau dynamic closely parallels the progressive neural alignment reported by Barczak et al. (2018).

**Figure 5.**
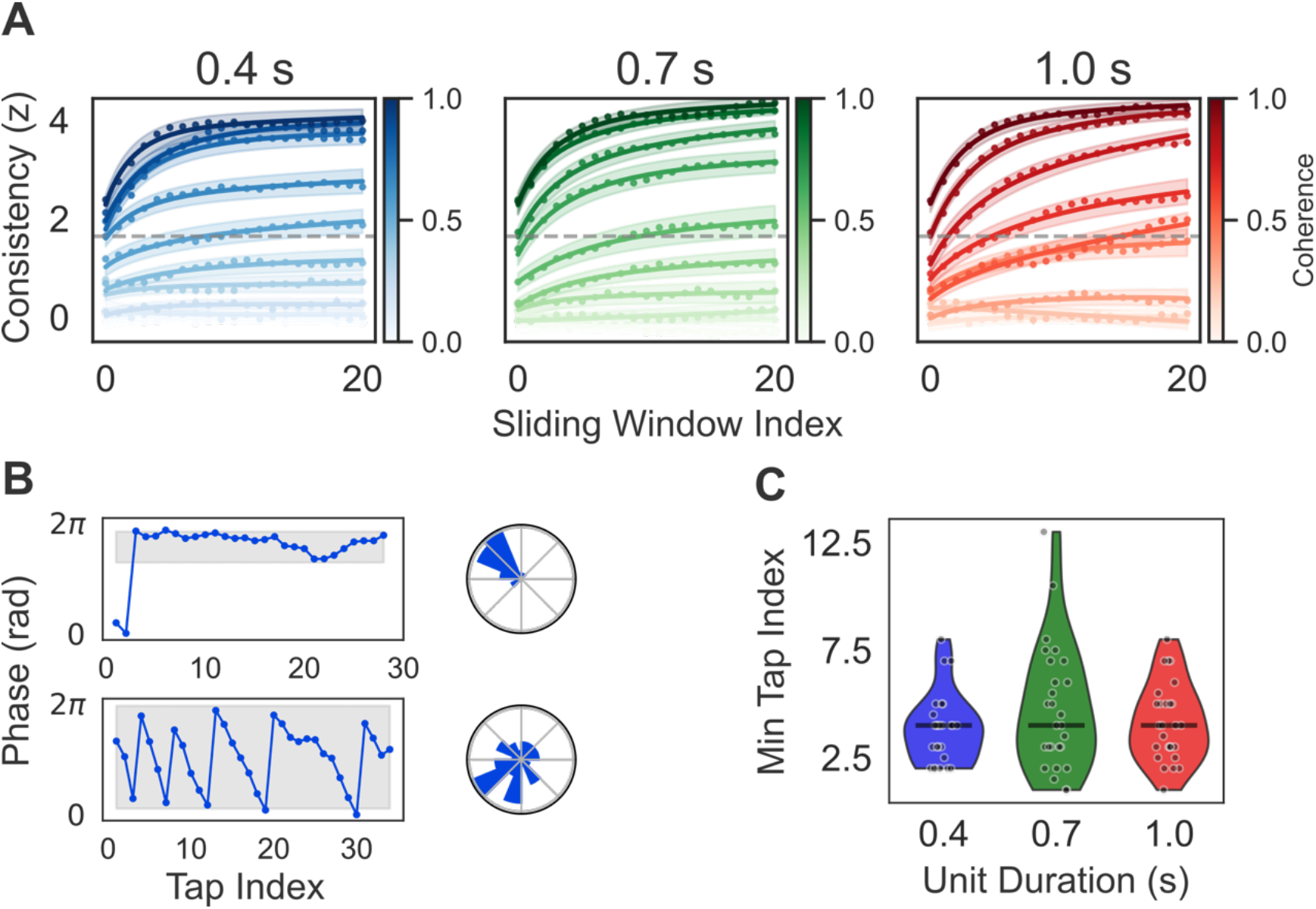
Temporal dynamics of evidence accumulation and maintenance in the SMS task. (A) Sliding-window analysis of tapping phase consistency in the sensorimotor synchronization (SMS) task. Tapping phase consistency, calculated from taps occurring within each window, is plotted as a function of sliding window index (see Materials and Methods). Each subplot corresponds to a different unit duration condition. Dots represent the mean z-scored resultant length (measure of tapping phase consistency) per window for each coherence level. Solid curves show the average saturating exponential fits across participants, with shaded regions indicating ±1 SEM. Repetition coherence levels are represented by the color gradient in the side color bar, where darker shades represent higher coherence and lighter shades indicate lower coherence. The dashed line marks the significance threshold (p = 0.05). (B) Examples of a trial with a narrow (upper) and broad (lower) stable phase range. Tapping points are shown on both linear and circular plots. In the linear plot, each dot represents the tapping phase relative to the repeating pattern onset. In the circular plot, the same tapping phases are displayed as a circular histogram. (C) Violin plot of the distribution of the earliest tap that aligns with the stable phase for the highest repetition coherence level, for all durations.

While the sliding-window analysis captures the overall synchronization dynamics, each window contains multiple taps. To gain a yet-more-fine-grained view, we analyzed individual taps in trials that showed successful synchronization, analogous to examining reaction times from correct trials. For each trial, we identified the earliest tap that aligned with the stable phase participants maintained in the final 20 cycles (see Materials and Methods; Figure 5B-C). This analysis was restricted to the highest coherence level, where the stable phase was sufficiently precise for the measure to be meaningful. At this coherence, taps converged around the fourth cycle, regardless of unit duration (Figure 5C) – the same cycle count, and the same scale-independence, observed in the detection task. This alignment in cycle count indicates that by the fourth cycle participants had formed a robust percept of the repeating pattern, sufficient for both detection and consistent tracking.

### A simple observer model reproduces each effect alone but not both together

To probe which aspects of the behavioral data follow from stimulus statistics and which require additional mechanisms, we implemented an autocorrelation-based observer model. The autocorrelation function (ACF) captures periodicity by comparing a signal with time-shifted versions of itself. It is a canonical framework for auditory periodicity extraction (de Cheveigné, 2003; Eck, 2006; Licklider, 1951; Pollack, 1978), with a biologically plausible counterpart in feature-detection circuits that retain information over short windows (Schöneich et al., 2015). Our stimuli lack perceptually resolvable individual tones and explicit broadband cues that mark boundaries of the repeating pattern. The ACF is well suited to such inputs as it can recover latent regularities even in their absence.

Rather than modeling specific auditory representations of the tone clouds, we used an abstract feature-level signal, allowing detectability to be examined independently of representational assumptions. A random base signal was tiled across cycles and mixed with independent noise according to a coherence parameter, at three signal lengths analogous to the short, medium and long unit durations in the experiment (see SI Materials and Methods). The temporal ACF of each signal was then computed. For the SMS task, ACF peak magnitude (Max Corr) was taken as a proxy for tapping phase consistency. For the detection task, a trial was registered as detected when the highest ACF peak exceeded a decision criterion (see SI Materials and Methods; Figure 6A).

**Figure 6.**
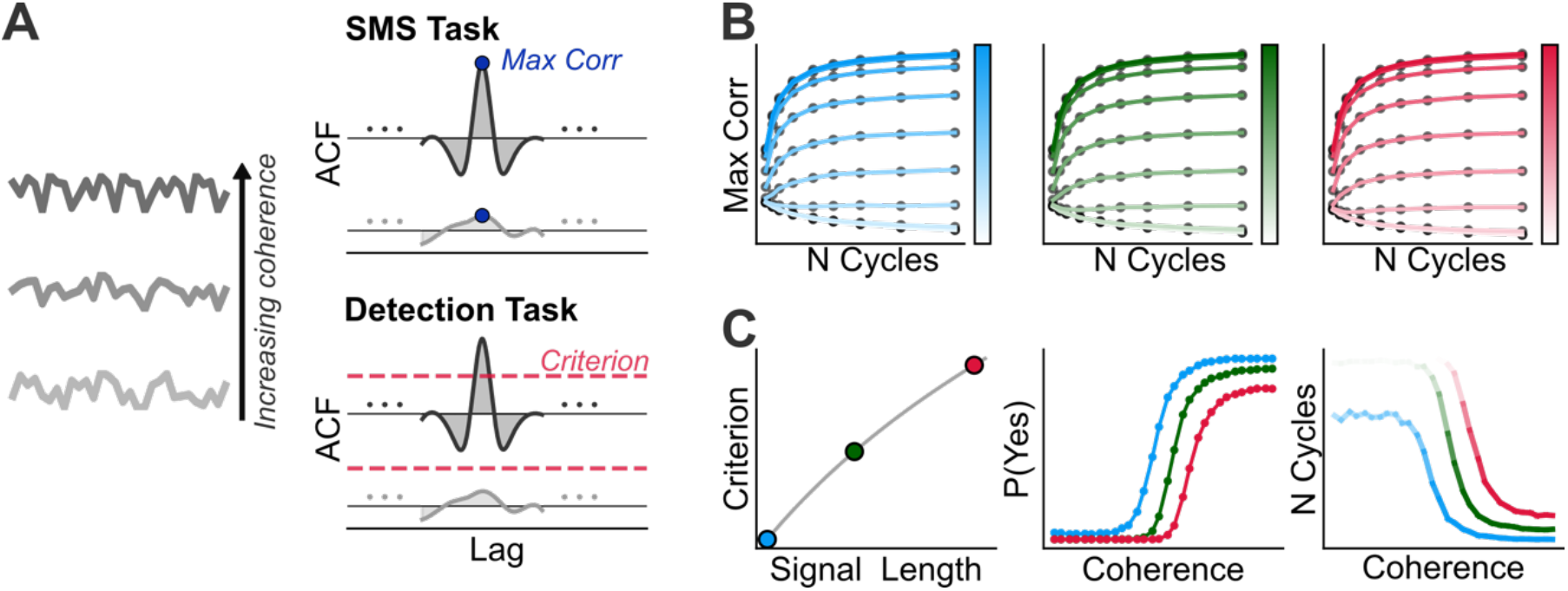
Observer model design and scale-dependent variant simulation results. (A) Model schematic. Left: example stimuli at increasing coherence levels. Right: the model computes the ACF of each input. For the SMS task, the maximum ACF peak magnitude (Max Corr) serves as a proxy for tapping phase consistency. For the detection task, repetition is registered when the ACF peak exceeds a decision criterion (dashed red line). SMS task simulations. Max Corr is plotted as a function of cycle number for each signal length (colors as in previous figures). Coherence levels are represented by the color gradient, with darker shades indicating higher coherence. (C) Detection task simulations with decision criterion scaled by signal length. Left: criterion as a function of signal length. Center: proportion of detected trials as a function of coherence, shown separately for each signal length (color coding as in B). Right: number of cycles required to reach criterion as a function of coherence for each signal length.

We first tested a scale-independent version with fixed internal parameters across signal lengths. The observer model reproduced the repetition-coherence-driven sigmoidal increase in detection performance and the buildup of tapping phase consistency across cycles. Both detection and SMS curves overlapped across signal lengths. The baseline model thus establishes that the coherence-driven effects in both tasks follow from stimulus statistics alone. Yet, it fails to capture the threshold shift in detection across unit durations, a shift absent in the SMS data (Figure S1).

Next, we tested a scale-dependent variant. We scaled the decision criterion with signal length, such that longer signals required stronger evidence to be classified as repeating (Figure 6C). The choice of stage was motivated by the distinct effect of unit duration on the two tasks. Detection requires a discrete decision, whereas SMS provides a continuous readout without an explicit decision stage. If the threshold shift arises at the decision stage rather than at the perceptual representation shared across tasks, then scaling the criterion should reproduce the dissociation.

This adjustment reproduced the asymmetry: distinct detection psychometric curves across signal lengths alongside scale-independent SMS performance (Figure 6B-C). A complementary variant scaling a representational rather than decisional parameter produced scale-dependent declines in both readouts, contrary to the human data (see SI Materials and Methods; Figure S2).

However, the scale-dependent variant exposes a tension that no parameter choice can resolve. The same manipulation that produces the detection threshold shift — a stricter criterion for longer signals — also predicts that longer signals should require more cycles to reach that criterion (Figure 6C). The human data show the opposite: at high coherence, the number of cycles required to reach detection is approximately the same across unit durations. This failure is structural rather than parametric. Any scale-dependent factor that makes detection harder for longer signals will necessarily require more integration to overcome it. Therefore, the scale-independent integration limit cannot coexist with the detection threshold shift within this framework. It points to an additional mechanism that bounds integration to a fixed number of cycles regardless of unit duration.

## Discussion

Using a paradigm designed to capture some of the complexity of real-world acoustic environments, we probed the parameters shaping auditory object formation and the minimal evidence required for object emergence. Listeners heard dense tone-cloud sequences containing no obvious segmentation cues, within which we embedded a frozen tone configuration. To succeed at detection, listeners had to organize this complex scene by identifying the frozen subset and grouping it into an auditory object. We first demonstrate that the segregation of the embedded object depends on the level of repetition coherence. Moreover, auditory objects emerge from the accumulation of sensory evidence across cycles, with the integration process constrained by intrinsic temporal windows. The resulting behavioral dynamics parallel known neural responses and provide a link between perception and physiology.

### Repetition Coherence

Our findings build on related work. McDermott et al. (2011) implemented a similar approach. They presented listeners with target-distractor mixtures whose statistics matched those of natural sounds. Within each trial, the target remained the same while the distractor changed on every presentation. They manipulated the number of presented mixtures to explore how repetition influenced signal decomposition. From a figure-ground perspective (Kubovy & Van Valkenburg, 2001), both tasks (theirs and ours) involve the challenge of grouping repeating elements into a coherent figure that can be segregated from ground. In vision, figure-ground segregation is argued to arise from competition for neural resources, with top-down mechanisms biasing perception toward specific objects (Desimone & Duncan, 1995; Lamme, 1995). To remove such generic top-down biases in our paradigm, we ‘conceptually reversed’ McDermott et al.’s (2011) approach. There were no predefined targets or distractors, nor any acoustic similarity with natural sounds. Participants encountered entirely novel sounds and constructed their own representation of a candidate figure, in real time, solely from bottom-up acoustic cues.

Not all repeating objects were detected by listeners. At low coherence, they failed to emerge, likely because informational masking from competing random tones prevented the pattern from being distinguished. Prior studies have demonstrated that immediate repetition of identical tones, chords, or brief tone bursts can overcome masking effects (Kidd et al., 1994, 2013; Micheyl et al., 2007; Teki et al., 2013). Our findings show that temporally extended repeating patterns do not yield the same benefit, which suggests that the temporal structure of repetition imposes constraints on the mechanisms that typically facilitate segregation.

The degree of repetition coherence not only affected whether listeners could segregate the repeating pattern but also shaped the temporal integration strategies they used. When there was no repeating pattern or when coherence was low, the integration strategy appeared to rely on absolute time. In contrast, at high coherence levels, the strategy appeared to shift to cycle-based integration. In the absolute-time strategy, waiting time was similar across unit durations. As a result, shorter units appeared to require more cycles than longer ones. By contrast, in all versions of the observer model, when coherence was low, longer units sampled more cycles than shorter ones to reach a decision. This discrepancy highlights a change in listeners’ temporal integration strategies depending on the available evidence, which our observer model does not account for.

The repetition coherence effects showed a dual pattern with respect to unit duration. In the repetition detection task, longer units were less likely to give rise to perceptual objects. On the other hand, a robust representation of an auditory object emerged after integrating the same number of cycles, independent of duration. This pattern indicates a dissociation: unit duration constrained the *likelihood* of object formation but not the integration process. Our observer model failed to reproduce this dissociation. In the scale-independent version, both the likelihood of detection and the number of cycles required were similar across unit durations. To account for differences in likelihood of object emergence, we scaled decision criteria with signal length. This scale-dependent version caused both detection likelihood and the required number of cycles to vary together. The human data, therefore, suggest the involvement of additional mechanisms not captured by the framework of our autocorrelation-based observer model.

One possible additional mechanism is motivated by the similarity of our data to Barczak et al.’s (2018) findings. They showed that neural alignment strength in A1 reached significance after the third presentation of a repeating segment. Although structurally more complex, our highest coherence condition nonetheless mirrors the critical aspect of their design: full repetition of a given tone sequence; and listeners detected the repetition and reached their stable tapping phase after the third presentation across all durations. This hints at a potentially rather direct perceptual-physiological correspondence and suggests that auditory object formation may be constrained by intrinsic temporal mechanisms. Specifically, phase alignment driven by repetition may require accumulation across multiple cycles and reach a level capable of triggering a behavioral outcome.

Converging evidence comes from Elhilali et al. (2009), who investigated how repeating target tones within complex masker sequences support detection. They observed improved behavioral detection with sufficient exposure, accompanied by a corresponding neural build-up (measured with MEG). Critically, this build-up was observable only when the analysis window spanned at least three rhythmic cycles of the target tone because the accumulation effect did not reflect a simple increase in power generated by individual targets. Rather, it arose from increasingly consistent phase alignment of neural responses to the target across successive cycles. Their findings suggest that auditory object formation relies on mechanisms that progressively refine the temporal alignment of neural activity across repeated exposures, thereby allowing stable perceptual representations to emerge.

Why does the likelihood of object formation decrease for longer durations? One possibility might be that auditory integration windows are bounded, rather than scaling with stimulus length (Giraud & Poeppel, 2012; Viemeister & Wakefield, 1991). This hypothesis might explain why shorter units yield higher detection accuracy: when an entire repeating pattern fits within a fixed integration window, evidence can accumulate fully and promote strong phase alignment in neural responses. In contrast, longer patterns may exceed these bounds or span multiple windows, which can prevent complete evidence accumulation. Consequently, the signal necessary to drive phase alignment, and ultimately trigger a behavioral outcome, fails to reach sufficient strength.

### Sensorimotor Synchronization

In the sensorimotor synchronization task, high repetition coherence produced a two-stage performance trajectory: an initial rapid improvement over the first few repetitions followed by a plateau. This build-up effect echoes classic reports from auditory stream segregation (Anstis & Saida, 1985; Bregman, 1978). Our findings also suggest, however, that the build-up does not reflect continuous accumulation, but instead a more categorical shift, from rapid evidence accumulation to a stabilized percept, beyond which additional exposure no longer influences behavior. This two-stage trajectory is consistent with the results of Barczak et al. (2018), who likewise reported a rapid rise in measured response followed by stabilization.

A finding specific to the SMS task is that unit duration no longer influenced performance; repetition coherence emerged as the primary determinant. One possible interpretation, informed by the simulations, is that the detection task might involve an additional decisional bottleneck, absent in SMS. This makes scale-dependent effects a by-product of decision criteria. This explanation alone is insufficient, though, as the same manipulation in the simulations also altered the number of cycles required for detection. These results suggest that further mechanisms contribute to the observed behavior patterns.

In summary, despite major differences in task demands, repetition coherence produced strikingly similar behavioral outcomes in both the likelihood of object formation and the stability with which that object could be tracked once formed. An autocorrelation-based observer model accounted for key coherence-driven effects across tasks, yet several important findings remained unexplained. These are: (1) the scale-independent nature of the temporal integration process, (2) the coherence-dependent shift in listeners’ integration strategies, and (3) the distinct influence of unit duration on detection versus synchronization. They all indicate the involvement of additional cognitive and neural mechanisms. Characterizing these mechanisms constitutes a critical next step toward a more comprehensive mechanistic account of how the auditory system structures the complex acoustic world.

## Materials and Methods

### Participants

Twenty-seven self-reported normal-hearing participants (18 females; mean age, 27.52; range, 21-39) completed two experimental sessions. Data from three additional participants were excluded due to excessive noise or technical problems with tapping signal acquisition. Participants provided written informed consent, approved by the Ethics Committee of the Medical Faculty of Goethe University, in accordance with the Declaration of Helsinki. They received monetary compensation.

### Stimuli: Tone Clouds

Stimuli were tone clouds consisting of brief pure-tone pips distributed pseudorandomly within time-frequency grids (Agus & Pressnitzer, 2021). Each grid cell contained one tone pip, with late-onset pips permitted to extend into the next cell. The lower and upper bounds of our tone-cloud units were 200 Hz and 3000 Hz. Consecutive frequencies differed by 0.4 octaves. This resulted in ten different frequency values constituting the rows of the grid structure.

The time step of each column was set to 0.05 s. The total number of columns depended on the length of a tone-cloud unit. There were three levels of unit duration: 0.4, 0.7, and 1.0 s. The duration of individual pips was 0.05 s, including 0.025-s onset-offset cosine ramps.

Each stimulus sequence was constructed by repeating a single tone-cloud unit 30 times. To create different levels of repetition coherence, we varied the proportion of tone pips within the unit that remained frozen across repetition cycles. We used ten values evenly spaced between 0 and 1.

### Experimental Tasks

There were two psychophysical tasks with identical conditions: a repetition detection task with a yes-no paradigm and a sensorimotor synchronization task. Separate stimulus sets with the same parameter values were used for the two tasks to prevent any learning effects.

#### Repetition Detection Task

Each block consisted of tone-cloud sequences that were either continuous (repetition coherence = 0; signal-absent) or repeating (nine non-zero repetition coherence levels; signal-present). Each block contained 108 trials (36 per unit duration condition). Participants completed seven blocks. Conditions were presented in a pseudorandom order. On each trial, participants listened to a tone-cloud sequence and indicated as quickly as possible whether they detected a repeating pattern. Although each sequence contained 30 repetitions, trials were terminated immediately upon response.

#### Sensorimotor Synchronization Task

Each block consisted of one trial per condition, resulting in 30 trials per block. Participants completed seven blocks. They were informed that sequences could be either continuous or repeating and were instructed to tap in synchrony with the perceived repeating pattern. Unlike the repetition detection task, participants heard the entire sequence containing 30 repetitions on each trial.

### Procedure

Participants took part in two separate experiments conducted on different days, with task order counterbalanced across individuals. They were seated in a soundproof booth, facing an LCD monitor for receiving instructions and feedback. Auditory stimuli were generated in Python (44.1 kHz, 16-bit) and delivered binaurally through electrodynamic headphones (Beyerdynamic DT770 PRO). The experiment was implemented using PsychoPy.

#### Repetition Detection Task

Participants first completed a brief training session to familiarize themselves with the stimuli and task structure (see SI Materials and Methods). Each experimental block began with a participant-initiated button press. Responses were collected via a response box (BBTK Response Pad), with “yes” and “no” mappings counterbalanced across participants. The inter-trial interval was 0.5 s. The full session lasted approximately two hours.

#### Sensorimotor Synchronization Task

The experiment began with a comparable training session (see SI Materials and Methods). Each experimental block was initiated by a participant’s button press. Trials began after a fixed 1 s delay. During each trial, participants tapped the index finger of their dominant hand on a desk next to a Schaller Oyster S/P contact microphone and continued tapping for the entire trial. The session lasted approximately two hours.

## Data Analysis

### Repetition Detection Task

#### Exclusion Criteria

Trials in which participants responded before hearing a full unit were excluded. RT outliers were identified separately for each repetition coherence and unit duration condition using the interquartile range (IQR) method (see SI Materials and Methods). Blocks with unusually low accuracy were excluded based on z-scores computed across blocks (|z| > 3). In total, 4.05% of trials were excluded.

#### Psychometric Function Fitting

Proportions of “yes” responses as a function of repetition coherence were modeled using a Weibull psychometric function, with threshold (#), slope ($), guess rate (%), and lapse rate (&) as free parameters. The parameters were estimated using maximum likelihood estimation under a binomial response model (see SI Materials and Methods).

#### Statistical Analysis

We conducted a 3 x 10 repeated-measures ANOVA to investigate the effects of repetition coherence and unit duration on detection performance, with participants’ average proportion of “yes” responses as the dependent variable. To assess the effect of unit duration on repetition detection threshold, we ran a separate one-way repeated-measures ANOVA using individual thresholds as the dependent variable.

To examine whether unit duration influenced RT and N cycles during successful repetition detection, trial-level data were summarized per participant, unit duration, and repetition coherence by computing median RT and N cycles for correct trials. Analyses were conducted separately for each repetition coherence level. Nonparametric within-subject Friedman tests assessed the effect of unit duration on RT and N cycles at each coherence level, followed by pairwise Wilcoxon signed-rank tests with Bonferroni correction.

### Sensorimotor Synchronization Task

#### Data Cleaning and Exclusion Criteria

Tapping data were recorded as audio. Individual taps were identified using an amplitude threshold applied to the waveform. Trials with one or fewer detected taps were excluded. Remaining trials were grouped by repetition coherence and unit duration, and outliers were removed using the IQR method. In total, 8.03% of trials were excluded.

#### Measure of Synchronization Success

For each trial, tap times in the last 20 repetition cycles were aligned to the corresponding cycle onset and converted into phase angles relative to the unit duration. Phase angles were represented as unit vectors, and the mean resultant vector was computed. The magnitude of the mean resultant vector, called the mean resultant length, reflects the concentration of tapping phases. To account for chance-level phase alignment, we simulated 1000 sets of random phases to generate a null distribution and computed a z-score relative to this distribution. This z-score served as our measure of tapping phase consistency (see SI Materials and Methods).

We also derived a binary outcome for each trial. A trial was classified as showing successful synchronization if its tapping phase consistency exceeded the 95^th^ percentile of the null distribution (p < 0.05). The proportion of successfully synchronized trials per condition was then used for psychometric function fitting.

#### Temporal Dynamics of Synchronization

To capture the temporal dynamics of synchronization performance, we applied a sliding window analysis. Each window spanned 10 repetition cycles and advanced by one cycle at each step. Within each window, we computed the tapping phase consistency. We then modeled the temporal evolution of tapping phase consistency using an exponential saturation function (see SI Materials and Methods).

We also examined individual taps within each trial. For each trial, we computed the circular mean and standard deviation of taps occurring within the last 20 repetition cycles, which represented the stable phase range tracked by participants within the repeating pattern. The earliest tap falling within this range was identified to determine when participants had locked onto the repeating pattern. Because this measure requires a precisely defined stable phase range, this analysis was restricted to the highest coherence level.

#### Psychometric Function Fitting

Overall synchronization performance across conditions was quantified by fitting a Weibull psychometric function (as in the detection task) to the binary successful synchronization measure. Parameters were estimated by minimizing the negative log-likelihood of a binomial response model.

#### Statistical Analysis

To examine how unit duration and repetition coherence influenced synchronization performance, we ran a 3 × 10 repeated-measures ANOVA with participants’ average proportion of successful synchronization trials as the dependent variable. To investigate the effect of unit duration on synchronization threshold, we conducted a separate one-way repeated-measures ANOVA using individual thresholds as the dependent variable.

## Observer Model

We developed an autocorrelation-based observer model to estimate latent temporal regularities. The model computes the ACF of the input signal, identifies peaks corresponding to repeating structures, and integrates information across cycles to stabilize periodicity estimates.

Synthetic signals were generated by repeating a base signal across multiple cycles and mixing it with independent noise to create varying repetition coherence levels. We tested three signal lengths, matched to the short, medium, and long unit durations used in the experiment. On each trial, internal (sensory) noise was added to the input signal to simulate trial-by-trial variability in the observer’s input. Internal noise was drawn from a uniform distribution on [-1, 1]. It was mean-centered and RMS-normalized before being mixed with the signal at a fixed level.

The temporal ACF of the noisy input was computed. We identified positive peaks and selected the largest non-zero-lag peak. The magnitude of that peak (denoted as Max Corr) served as the model’s readout of periodic structure. For the SMS task, Max Corr indexed tapping phase consistency on a continuous scale. For the detection task, a trial counted as detected when Max Corr exceeded a decision criterion. To estimate the number of cycles required for detection, signals were grown one cycle at a time and Max Corr recomputed at each step. We then recorded the cycle at which Max Corr first exceeded the criterion.

We tested two main variants. In the scale-independent variant, internal noise and decision criterion were held constant across signal lengths. In the scale-dependent (criterion scaling) variant, the criterion increased with signal length according to a normalized logarithmic function, such that longer signals required higher Max Corr to cross the decision criterion. We also added small trial-level Gaussian noise to the criterion to capture within-observer variability. A complementary variant in which internal noise scaled with signal length is reported in the SI. Each condition was simulated across multiple runs. Full implementation details, parameters, and simulation procedures are provided in SI Materials and Methods.

## Supporting information

Materials and Methods, Results

## Acknowledgments

This work was supported by Max Planck School of Cognition and the Ernst Struengmann Institute. For helpful feedback and comments, we would like to thank Vani Rajendran, Daniel Pressnitzer, Jiajie Zou, Leo Zeine, and all members of the Poeppel lab.

## Author Contributions

**Conceptualization:** Berfin Bastug, Yue Sun, Federico Adolfi, Erich Schröger, David Poeppel.

**Formal analysis:** Berfin Bastug.

**Investigation:** Berfin Bastug.

**Methodology:** Berfin Bastug, Yue Sun, Federico Adolfi.

**Supervision:** David Poeppel, Erich Schröger.

**Visualization:** Berfin Bastug.

**Writing – original draft:** Berfin Bastug, Yue Sun, Federico Adolfi, Erich Schröger, David Poeppel.

**Writing – review & editing:** Berfin Bastug, Yue Sun, Federico Adolfi, Erich Schröger, David Poeppel.

