## Supplementary material for "Computational Principles of Auditory Object Formation Are Revealed by Repetition Coherence": Materials and Methods, Results

### Supplemental Information

### Materials and Methods

#### Training Procedure

*Repetition Detection Task*

Participants listened to sequences in which the tone clouds of three different unit durations repeated ten times. Then, they completed a short training block. This block consisted of 10 trials drawn from a representative subset of the experimental conditions. Half of the trials were signal-absent, while the other half were signal-present trials with repetition coherence levels of 0.2, 0.7 and 1. The training block was repeated, if necessary, until participants achieved at least 75% accuracy.

*Sensorimotor Synchronization Task*

Participants were introduced to fully repeating tone cloud sequences and the range of unit durations used in the experiment. After passively listening to these sequences, they heard another set of fully repeating stimuli and were asked to tap in synchrony with the perceived repeating pattern to ensure they understood the task requirement.

#### Data Analysis

*Repetition Detection Task*

*Exclusion Criteria:* RT outliers were identified separately for each combination of repetition coherence and unit duration condition using the interquartile range (IQR) method, based on the 25th (Q1) and 75th (Q3) percentiles. Trials with RTs outside the range of $Q_1 - 1.5 \times \mathrm{IQR}$ to $Q_3 + 1.5 \times \mathrm{IQR}$ were excluded.

*Psychometric Function Fitting:* We used a Weibull function to model the relationship between the proportion of “yes” responses and repetition coherence:

$$\psi(x) = \gamma + (1 - \gamma - \lambda)\left[1 - e^{-(x/\alpha)^{\beta}}\right]$$

where $x$ denotes repetition coherence, $\alpha$ the detection threshold, $\beta$ the slope of the function, $\gamma$ the guess rate, and $\lambda$ the lapse rate.

Parameters were estimated via maximum likelihood estimation (MLE) by minimizing the negative log-likelihood (NLL) of a binomial response model. At each coherence level $i$, the number of "yes" responses $y_i$ was modeled as a binomial draw based on the predicted probability $p_i$ from the psychometric function and the number of trials $n_i$.

$$NLL = -\sum_i [y_i \log(p_i) + (n_i - y_i)\log(1 - p_i)]$$

*Sensorimotor Synchronization Task*

*Measure of Synchronization Success:* For each trial, we used the tap onset times from the last 20 repetition cycles. Each tap onset time $t_i$ was aligned to the most recent cycle onset $c_i$, $\delta_i = t_i - c_i$. These alignments were converted into phase angles, $\theta_i = (2\pi\delta_i)\,/\,T$, by normalizing with respect to the unit duration $T$. Phase angles were represented as complex unit vectors $\upsilon_i = \cos(\theta_i) + i\ \sin(\theta_i)$, and the mean resultant vector was computed.

$$\bar{R} = \frac{1}{n}\sum_{i=1}^{n} \upsilon_i$$

Then, we calculated its magnitude $r = |\bar{R}|$, which is called mean resultant length. We simulated 1000 sets of n random phases uniformly sampled from $[0,2\pi]$, and computed their mean resultant vector lengths. From these, we derived the mean $(\mu_{\text{null}})$ and standard deviation $(\sigma_{\text{null}})$. We calculated the z-score to account for chance-level phase alignment, which we used as a measure of tapping phase consistency.

$$z = \frac{r_{\text{obs}} - \mu_{\text{null}}}{\sigma_{\text{null}}}$$

A trial was classified as successfully synchronized if its z-score exceeded the 95$^{\text{th}}$ percentile of the null distribution ($p < 0.05$). We then used the proportion of successfully synchronized trials per condition for psychometric function fitting. Parameters were estimated using the same Weibull function and negative log-likelihood minimization procedure described above for the repetition detection task.

*Temporal Dynamics of Synchronization:* We characterized the dynamics of tapping phase consistency with an exponential saturation function: $f(x) = a(1 - e^{-bx}) + c$, where $a$ is the asymptotic (maximum) level of phase consistency, $b$ is the rate at which phase consistency stabilizes, and $c$ is the baseline phase consistency.

**Observer Model**

A base signal was generated as a random one-dimensional vector of length $T$. Signal lengths were drawn from a logarithmically spaced domain ranging from 20 to 40 samples. Stimuli consisting of $N$ repetition cycles were constructed by concatenating this base signal $N$ times.

To manipulate repetition coherence, an independent noise signal of equal length was generated and linearly mixed with the repeating base signal. Repetition coherence $c \in [0,1]$ controlled the relative contributions of the repeating and noise signal components:

$$x(t) = c \cdot x_{\text{repeat}}(t) + (1 - c) \cdot x_{\text{noise}}(t)$$

where $x_{\text{repeat}}(t)$ is the repeating base signal and $x_{\text{noise}}$ is an independent noise signal. Coherence values were sampled from 30 linearly spaced levels between 0 and 1.

All resulting signals were mean-centered and root-mean-square (RMS)-normalized to ensure that differences in autocorrelation magnitude were driven by repetition coherence rather than overall signal energy.

An additional internal (sensory) noise term was added to each stimulus prior to temporal autocorrelation function (ACF) computation. Internal noise was drawn from a uniform distribution over [-1, 1], mean-centered, and RMS-normalized. This internal noise was then weighted by a gain parameter and added to the stimulus. Unless otherwise specified, the internal noise gain was fixed at 0.15 across all signal lengths.

The observer computed the ACF of the entire signal. The magnitude of the largest positive peak (excluding the zero-lag peak), denoted Max Corr, served as the model's readout of periodic structure.

*Output Mapping*

*Repetition Detection Task:* Repetition detection was modeled as a criterion-crossing process. A stimulus was classified as containing a repeating pattern when Max Corr exceeded a decision criterion. Decision criteria were sampled from 10 linearly spaced values between 0.3 and 0.4, spanning conservative to liberal observers. Model outputs were aggregated across criteria to characterize performance trends independent of a specific criterion setting.

The observer evaluated signals incrementally as the repetition cycles accumulated from 1 to 30. Detection time was defined as the first cycle at which Max Corr exceeded the criterion. If the criterion was not reached within the maximum number of cycles, the stimulus was classified as undetected.

*Sensorimotor Synchronization Task*: For the sensorimotor synchronization (SMS) task, Max Corr served as a continuous proxy for tapping phase consistency. Larger Max Corr values correspond to stronger periodic structure in the signal. We therefore assumed that this stronger periodicity would in turn produce more concentrated tapping phases around a stable position.

For all model variants, Max Corr and detection decisions were recorded on every trial and aggregated across runs for analysis.

*Model Comparisons with Different Variants*

*Scale-Independent Model with Fixed Internal Parameters:* The decision criteria and noise gain described above were applied uniformly across signal lengths. Small trial-level Gaussian noise ($\sigma = 0.07$) was added to the criterion on each trial to capture within-observer variability. Each condition was simulated over 50 independent runs per base criterion (10 base criteria x 50 runs = 500 total samples per condition).

*Scale-Dependent Model with Increasing Decision Criterion with Signal Length:* In this variant, sensitivity to repetition was modulated at the decision stage by allowing the decision criterion to increase logarithmically with signal length. For each simulated trial, the effective decision criterion was computed as:

$$c = c_{\text{base}} + \frac{log(L/L_{min})}{log(L_{max}/L_{min})} \cdot s$$

where $C_{base}$is the base criterion, $L$ is the signal length, $L_{min}$ and $L_{max}$ are the shortest and longest signal lengths, and $s$ is the slope of the logarithmic increase. As in the scale-independent variant, small trial-level Gaussian noise was added to the criterion. Effective criterion values exceeding 1 were clipped. Each condition was simulated over 50 independent runs per base criterion (500 total samples per condition).

*Scale-Dependent Model with Increasing Internal Noise with Signal Length:* This time, the internal (sensory) noise gain increased logarithmically with signal length, while the decision criterion was not scaled with signal length. The effective noise gain for each trial was computed as:

$$n = n_{\text{base}} + \frac{log(L/L_{min})}{log(L_{max}/L_{min})} \cdot s$$

where $n_{base}$ is the base noise gain and $s$ is the slope of the logarithmic increase. Decision criteria were drawn from the same range as in the previous variants, with trial-level Gaussian noise. Each condition was simulated over 50 independent runs per base criterion (500 total samples per condition).

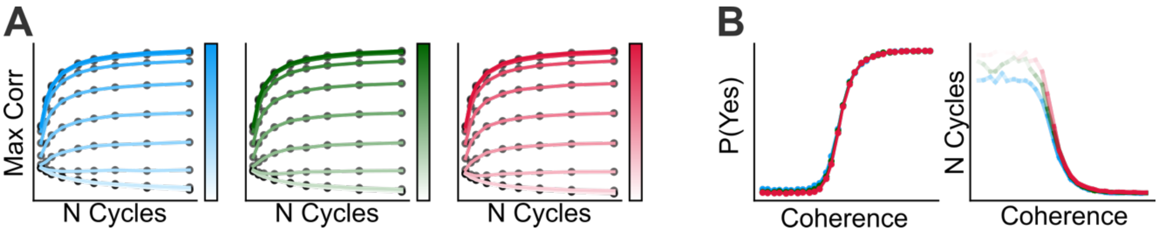


**Figure S1.** Simulation results of the scale-independent model. (A) SMS task simulations. Max Corr is plotted as a function of cycle number for each signal length (short: blue, medium: green, long: red). Coherence levels are represented by the color gradient, with darker shades indicating higher coherence. (B) Detection task simulations. Left: proportion of detected trials as a function of coherence, shown separately for each signal length (color coding as in A; curves largely overlap). Right: number of cycles required to reach criterion as a function of coherence for each signal length.

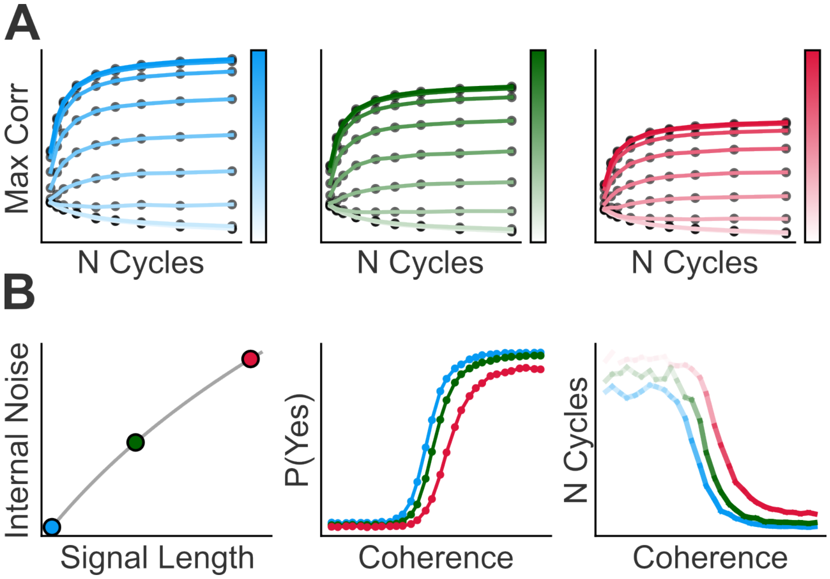

**Figure S2.** Simulation results of the scale-dependent model with internal noise scaled by signal length. (A) SMS task simulations. Max Corr is plotted as a function of cycle number for each signal length (short: blue, medium: green, long: red) and coherence level (darker shades indicate higher coherence). (B) Detection task simulations. Left: internal noise gain as a function of signal length. Center: proportion of detected trials as a function of coherence for each signal length (color coding as in A). Right: number of cycles required to reach criterion, plotted against coherence for each signal length.

**Table S1.** Mean proportion yes and proportion of successful synchronization trials across unit durations

| Repetition Detection | | | | | | |
|---|---|---|---|---|---|---|
| **Measure** | **0.4-s** | | **0.7-s** | | **1-s** | |
| | ***M*** | ***SD*** | ***M*** | ***SD*** | ***M*** | ***SD*** |
| Prop Correct | 0.68 | 0.1 | 0.56 | 0.12 | 0.45 | 0.15 |

| Sensorimotor Synchronization | | | | | | |
|---|---|---|---|---|---|---|
| **Measure** | **0.4-s** | | **0.7-s** | | **1-s** | |
| | ***M*** | ***SD*** | ***M*** | ***SD*** | ***M*** | ***SD*** |
| Prop Successful | 0.57 | 0.13 | 0.55 | 0.12 | 0.54 | 0.1 |

**Table S2.** Repetition coherence thresholds (50%) and repeated-measures ANOVA results for repetition detection and sensorimotor synchronization tasks.

**Repetition Detection**

| **Measure** | **0.4-s** | | **0.7-s** | | **1-s** | | **F (1.56, 40.60)** | $\eta_G^2$ |
|---|---|---|---|---|---|---|---|---|
| | ***M*** | ***SD*** | ***M*** | ***SD*** | ***M*** | ***SD*** | | |
| Threshold | 0.31 | 0.12 | 0.47 | 0.13 | 0.58 | 0.19 | 96.96 ** | 0.36 |

**Sensorimotor Synchronization**

| **Measure** | **0.4-s** | | **0.7-s** | | **1-s** | | **F (2, 52)** | $\eta_G^2$ |
|---|---|---|---|---|---|---|---|---|
| | ***M*** | ***SD*** | ***M*** | ***SD*** | ***M*** | ***SD*** | | |
| Threshold | 0.49 | 0.16 | 0.51 | 0.13 | 0.49 | 0.13 | 0.44 | 0.006 |

Degrees of freedom and *p*-values were corrected using the Greenhouse–Geisser procedure where sphericity was violated. Effect sizes are reported as generalized eta squared ($\eta_G^2$). *p < .05, **p < .01.

**Table S3.** RT and N cycles as a function of repetition coherence and unit duration

| Coh. | Measure | 0.4-s (M, SD) | 0.7-s (M, SD) | 1-s (M, SD) | $\chi^2(2)$ | *p* | Post-hoc summary |
|---|---|---|---|---|---|---|---|
| 0 | RT | 5.02, 1.18 | 5.07, 1.22 | 4.97, 1.14 | 0.52 | 0.77 | - |
| | $N_{cycles}$ | 12.5, 2.94 | 7.25, 1.74 | 4.97, 1.14 | 54 | <.01 | 1 < 0.7<br>0.7 < 0.4<br>1 < 0.4 |
| 0.11 | RT | 4.67, 1.77 | 5.16, 2.64 | 5.91, 2.19 | 6 | .05 | 0.4 < 1 |
| | $N_{cycles}$ | 11.7, 4.42 | 7.38, 3.77 | 5.91, 2.19 | 14 | < .01 | 0.7 < 0.4<br>1 < 0.4 |
| 0.22 | RT | 4.22, 1.56 | 4.53, 1.65 | 4.53, 1.72 | 3.29 | .19 | - |
| | $N_{cycles}$ | 10.5, 3.90 | 6.47, 2.35 | 4.53, 1.72 | 23.06 | < .01 | 1 < 0.7<br>0.7 < 0.4<br>1 < 0.4 |
| 0.33 | RT | 4.04, 1.45 | 5.04, 1.85 | 5.42, 1.83 | 11.37 | < .01 | 0.4 < 0.7<br>0.4 < 1 |
| | $N_{cycles}$ | 10.1, 3.61 | 7.20, 2.64 | 5.42, 1.83 | 23.47 | < .01 | 0.7 < 0.4<br>1 < 0.4 |
| 0.44 | RT | 3.45, 1.45 | 5.39, 2.23 | 5.31, 2.36 | 18.87 | < .01 | 0.4 < 0.7<br>0.4 < 1 |
| | $N_{cycles}$ | 8.63, 3.63 | 7.69, 3.19 | 5.31, 2.36 | 20.96 | < .01 | 1 < 0.4<br>1 < 0.7 |
| 0.56 | RT | 3.00, 0.90 | 4.52, 1.57 | 5.82, 1.93 | 40.56 | < .01 | 0.4 < 0.7<br>0.4 < 1<br>0.7 < 1 |

| | | | | | | | |
|---|---|---|---|---|---|---|---|
| | $N_{cycles}$ | 7.50, 2.25 | 6.45, 2.24 | 5.82, 1.93 | 23.28 | < .01 | 0.7 < 0.4<br>1 < 0.4 |
| 0.67 | RT | 2.33, 0.70 | 4.04, 1.35 | 5.80, 1.93 | 52.07 | < .01 | 0.4 < 0.7<br>0.4 < 1<br>0.7 < 1 |
| | $N_{cycles}$ | 5.84, 1.76 | 5.76, 1.93 | 5.80, 1.93 | 0.52 | .77 | - |
| 0.78 | RT | 1.96, 0.40 | 3.68, 1.01 | 5.35, 1.65 | 54 | < .01 | 0.4 < 0.7<br>0.4 < 1<br>0.7 < 1 |
| | $N_{cycles}$ | 4.89, 0.99 | 5.25, 1.44 | 5.35, 1.65 | 3.19 | .2 | - |
| 0.89 | RT | 1.84, 0.32 | 3.09, 0.69 | 4.52, 1.24 | 52.07 | < .01 | 0.4 < 0.7<br>0.4 < 1<br>0.7 < 1 |
| | $N_{cycles}$ | 4.60, 0.80 | 4.42, 0.98 | 4.52, 1.24 | 3.19 | .20 | - |
| 1 | RT | 1.74, 0.25 | 2.74, 0.57 | 4.17, 1.14 | 54 | < .01 | 0.4 < 0.7<br>0.4 < 1<br>0.7 < 1 |
| | $N_{cycles}$ | 4.36, 0.63 | 3.91, 0.81 | 4.17, 1.14 | 10.67 | < .01 | 0.7 < 0.4 |

Values are means of participant-level medians. Differences across unit durations at each coherence level were assessed using one-way Friedman tests, followed by pairwise Wilcoxon signed-rank tests with Bonferroni correction where applicable.

**Table S4.** Effects of repetition coherence and unit duration on repetition detection and sensorimotor synchronization performance

| **Effect** | **Repetition Detection** | | | **Sensorimotor Synchronization** | | |
|---|---|---|---|---|---|---|
| | $df$ | $F$ | $\eta_G^2$ | $df$ | $F$ | $\eta_G^2$ |
| Unit Duration | 1.59, 41.43 | 186.05** | 0.23 | 2, 52 | 0.4 | 0.002 |
| Coherence | 2.29, 59.60 | 352.05** | 0.77 | 3.53, 91.89 | 247.69** | 0.7 |
| Unit Dur x Coh | 6.70, 174.15 | 22.94** | 0.16 | 18, 468 | 1.57 | 0.02 |

Results of 3 × 10 repeated-measures ANOVAs with unit duration and repetition coherence as within-subjects factors. The dependent variable was the proportion of "yes" responses for repetition detection and the proportion of successfully synchronized trials for sensorimotor synchronization. Degrees of freedom and *p*-values were Greenhouse–Geisser corrected where sphericity was violated. Effect sizes are reported as generalized eta squared ($\eta_G^2$). *p < .05, **p < .01.
